# MQ: Tool for HR-AM Based Quantitative Analysis for Direct MS Workflow

**DOI:** 10.1101/2020.01.19.912055

**Authors:** Avinash Ghanate, Dharmesh Parmar, Nivedita Bhattacharya, Ijaz Inamdar, Gayatri Phadke, Venkateswarlu Panchagnula

## Abstract

There is an increased interest in using high-resolution mass spectrometry (HRMS) in the full scan mode (FS) for simultaneous qualitative and quantitative analysis with accurate mass based extracted ion chromatograms (HRMS-AM) for metabolomic profiling workflows. Herein, the capabilities and features of ‘MQ’, a common platform developed to support quantitative analysis from HRMS data are described. Module for qualitative annotation utilizes peak features that include mass accuracy, peak width, and relative abundance of isotopic distribution that offer reliable metabolic confirmation. Additionally, MQ provides multivariate clustering with Principal Component Analysis and a module for high-throughput database query based on user configurable annotated metabolite lists. MQ is capable of handling system-agnostic data formats from both direct HRMS and LC-HRMS analysis. Data processed using MQ for LC-HRMS-AM quantitation of amino acids benchmarked using a commercial tool is presented that validates the quantitative capabilities of the algorithm developed. Datasets representative of three different HRMS-AM based analysis scenarios have been showcased to demonstrate the utility of MQ.

**Availability and implementation:** Freely available on web at, http://www.ldims.com/services/software. MQ is implemented in Java and supported on Linux and Microsoft Windows system having Java runtime environment (JRE) pre-installed.

## 1 Introduction

Contemporary high-resolution mass spectrometry (HRMS) provide millidalton (mDa) level resolution and mass-to-charge ratio (*m/z*) measurement within a few parts per million (ppm) accuracy of the exact mass as opposed to nominal mass based analysis on conventional low resolution mass spectrometry (MS) platforms. It is now theoretically possible to accurately profile and qualify a few thousands of distinct metabolites from diverse biological sources within a single HRMS scan. HRMS coupled to a chromatographic front end (gas or liquid chromatography (GC or LC)) is usually the preferred choice for metabolic profiling.[21,6] Various chromatography-free direct and ambient ionization methods can be used subsequently, in conjunction with HRMS analyzers to further aid high-throughput profiling, occasionally even directly from sample surfaces.[23] Mass spectral imaging (MSI) using direct ionization methods has shown potential to obtain mechanistic and molecular insights at a cellular, organ and systemic level.[13]

Analysis of HRMS data for diverse applications requires efficient algorithmic tools to extract and process relevant information from raw data. A common data analysis pipeline followed for processing HRMS data is represented in Figure 1. Various proprietary and a few open source tools that can support HRMS data analysis are currently available.[18,19,10,17,3,20] (Refer Table 1 for brief list of available tools and supported platform) Usually proprietary softwares for data analysis come bundled with mass spectrometry instruments and support only specific file formats. Inherent raw data inflexibilities can at times be limiting especially if one were to work across platforms from different manufacturers. A few open-source tools such as mMass [18,19], MZmine [10], XCMS [17], Mascot [3] and TOPP [20] are also available. Some of these tools do not include options for absolute quantitation and are geared towards particular workflows (for example proteomics, metabolomics, and lipidomics profiling). Also, data handling in the majority of data processing tools for quantitative analysis is pegged with chromatography-based workflows leaving little room for those exploring direct HRMS measurements devoid of chromatography. Most of the available open source tools were developed using Python, C++ or R platform depending upon different features offered such as the availability of numerical or statistical libraries and modular structure making it flexible to integrate in third party applications. Availability of Java based open source tools for MS data analysis is limited although the Java platform is powered with many more features and has an array of libraries for numerical or multivariate analysis along with recent active development of Mass Spectral Development Kit (MSDK) (https://msdk.github.io/).

**Table 1.** List of available open-source software tools or algorithms for data analysis enlisting offered features for data processing

| Tools/<br>Algorithm | Data<br>type | Data pre-processing |  |  |  | Feature analysis |  |  |
| --- | --- | --- | --- | --- | --- | --- | --- | --- |
|  |  | S/D | Base | Norm | Align | FX | Annot | Quant |
| MQ | NC/LC<br>MS |  |  | ✓ | NA | ✓ | ✓ | ✓ |
| PROcess | NC MS |  | ✓ | ✓ |  | ✓ |  |  |
| Cromwell | NC MS |  | ✓ | ✓ |  | ✓ |  |  |
| SpecAlign | NC MS | ✓ | ✓ |  | ✓ | ✓ |  |  |
| MassSpecWavelet | NC MS |  |  |  |  | ✓ |  |  |
| FlavonQ | LC MS |  |  |  |  | ✓ | ✓ | ✓ |
| MS-DIAL | LC<br>MS/MS |  |  |  |  | ✓ |  | ✓ |
| Met-XAlign | LC MS |  |  |  | ✓ |  |  |  |
| MetCOFEA | LC MS |  |  |  | ✓ |  |  |  |
| Open-SWATH | LC<br>MS/MS |  |  |  |  |  | ✓ |  |
| MS2Analyzer | NC<br>MS/MS |  |  |  |  |  | ✓ |  |
| TIPick | LC MS | ✓ |  |  |  | ✓ |  |  |
| MultiAlign | LC MS |  |  |  | ✓ |  |  |  |
| xMSanalyzer | LC MS |  |  |  |  | ✓ | ✓ |  |
| MetSign |  |  |  | ✓ |  | ✓ |  |  |
| SIMA | LC MS |  |  |  | ✓ |  |  |  |
| MSClust | LC/GC<br>MS |  |  |  | ✓ |  |  |  |
| PeakMl/MZmatch | LC MS |  |  |  |  | ✓ | ✓ |  |

Note: <sup>NC</sup> Direct MS, <sup>LC</sup> LCMS, <sup>GC</sup> GCMS, <sup>S/D</sup> Smoothing/Denoise, <sup>Base</sup> Baseline correction, <sup>Norm</sup> Normalization, <sup>Align</sup> Alignment, <sup>FX</sup> Feature extraction, <sup>Annot</sup> Annotation, <sup>Quant</sup> Quantitation

**Fig. 1.**
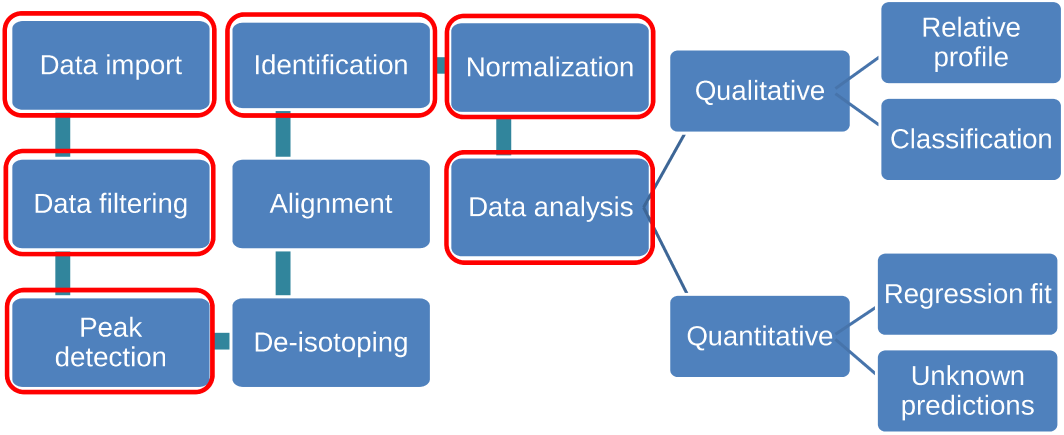
Detailed workflow for HRMS Data analysis. Highlighted aspects are featured in MQ

Herein, we report ‘MQ’, an algorithm developed using Java platform that attempts to support high throughput HRMS-based targeted metabolite quantitation workflows subsequent to global metabolomic profiling and qualification. MQ has been developed in our group with a broad perspective to aid qualitative and quantitative HRMS data analysis for direct MS analysis, especially to support MALDI-MS workflows. The algorithm has been continuously improvised, rigorously tested and benchmarked vis-à-vis proprietary Xcalibur™(Thermo Scientific) software that is code of federal regulations compliant (CFR, FDA). Herein, we describe various features of MQ and showcase a comparative study for the assessment of its quantitative performance in direct ionization and LC-HRMS based metabolic analysis for a targeted set of analytes. Preliminary versions of MQ were previously used in quantitative non-chromatographic laser desorption ionization mass spectrometry-based analysis workflows [2,16].

## 2 Materials and Methods

### 2.1 Chemicals

LCMS grade methanol and acetonitrile was procured from J.T.Baker (India). S-adenosylmethionine (SAM), S-adenosine-L-homocysteine (SAH), trifluoroacetic acid (TFA), verapamil ((RS)-2- (3,4-dimethoxyphenyl)-5-2-prop-2-ylpentanenitrile) and 2, 5-dihydroxy benzoic acid (2, 5-DHB) were purchased from Sigma-Aldrich. Melamine (2, 4, 6-triamino-1, 3, 5-triazine) was purchased from Loba Chemie (India). Atorvastatin ((3R,5R)-7-[2-(4-Fluorophenyl)-3-phenyl-4-(phenylcarbamoyl)-5-propan-2-ylpyrrol-1-yl]-3,5-dihydroxyheptanoic acid) was obtained as gratis sample from Mylan Laboratories limited, Hyderabad, India. Deionized water with specific resistance 18.2 M*Ω* cm^-1^ was obtained from Milli-Q unit (Merck Millipore). Mammalian cell culture medium composed of DMEM (Sigma-Aldrich, D6046) supplemented with MEM Non-essential amino acids solution (Sigma-Aldrich, M7145), was used as standard mixture for all amino acids. All the chemicals used in this study were of analytical grade.

### 2.2 LC-HRMS based quantitative analysis of amino acids

The LC-HRMS instrumentation consisted of autosampler (Accela Open Autosampler, Thermo Scientific) and liquid chromatograph (Accela 1250, Thermo Scientific) in tandem with the Q-Exactive (Thermo Scientific) high resolution mass spectrometer equipped with a heated electrospray ionization (HESI) interface. Instrument operation and data acquisition was performed using the ‘Xcalibur™’ platform software (Thermo Scientific). A C18 Hypersil gold column (10 cm x 2.1 mm x 3.0 *µ*M) by Thermo Scientific was used for eluting the samples prior to the ESI. The mass analyzer was operated in positive ion mode and data was acquired in triplicates within a mass range of 60-900 *m/z* at 70,000 FWHM resolution.

For quantitative analysis, mammalian cell culture medium and extracellular sample extracts for two neuronal cancer cell lines (U87MG and NSP) that were characterized in a parallel published study[4], were utilized. Cell culture medium standard mixture was serially diluted to generate calibration curves for the ranges reported in Table 3. A total of 10 calibration levels and 2 quality control (QC) samples were used. To investigate differential metabolic exchange profile, sample extracts from every 24 hr were pooled across cellular culture growth over 7 days. A 100 *µ*L of such sample extract was mixed with 400 *µ*L of chilled methanol (previously stored in -80C). The solution was thoroughly mixed for 2 mins followed by centrifugation for 15 mins at 5000 rpm (4C). The tubes were carefully removed, 300 *µ*L of supernatant was withdrawn and transferred into a fresh tube. A two-step serial dilution of supernatant was performed using 50 % acetonitrile in water. In the first step, 50 *µ*L of supernatant was thoroughly mixed with 450 *µ*L of diluent. This solution was further diluted by mixing 100 *µ*L of sample solution with 400 *µ*L of diluent. These samples along with standard mix were uniformly spiked with the 2 *µ*M solution of verapamil as internal standard to evaluate the performance and for data normalization. The solutions were thoroughly mixed and were then analyzed on LC-HRMS system. Following the acquisition, the raw data was analyzed using proprietary ‘Quan-browser’ module from Xcalibur™. Separately, data analysis using MQ was carried out as outlined below.

**Table 2.** Statistical evaluation of quantitative features preserved post data transformation into two dimensional mass spectral profile

| | Mean peak<br>area<br>(MQ) | Mean peak<br>area<br>(Xcalibur<sup>TM</sup>) | Pearson<br>correlation($r^2$) | F-test<br>statistics | F-test<br>P value |
| --- | --- | --- | --- | --- | --- |
| Ala | 7.90E+05 | 2.30E+07 | 0.9900 | 9.17E-11 | 1.83E-10 |
| Arg | 2.73E+06 | 1.87E+08 | 0.9974 | 2.47E-09 | 4.94E-09 |
| Asn | 2.39E+05 | 1.67E+07 | 0.9678 | 1.45E-09 | 2.90E-09 |
| Asp | 4.49E+05 | 3.22E+07 | 0.7993 | 4.42E-11 | 8.83E-11 |
| Cyst | 4.23E+05 | 2.90E+07 | 0.9999 | 3.94E-09 | 7.87E-09 |
| Glu | 4.08E+05 | 2.77E+07 | 0.9894 | 1.27E-09 | 2.55E-09 |
| Gln | 8.40E+06 | 5.70E+08 | 0.997 | 1.12E-09 | 2.24E-09 |
| Gly | 1.15E+06 | 7.31E+07 | 0.9995 | 1.97E-10 | 3.93E-10 |
| His | 1.33E+06 | 9.16E+07 | 0.9997 | 1.50E-09 | 2.99E-09 |
| Leu | 4.06E+07 | 2.78E+09 | 0.9815 | 8.11E-10 | 1.62E-09 |
| LYS | 4.31E+06 | 2.95E+08 | 0.9999 | 1.05E-09 | 2.10E-09 |
| MET | 1.97E+06 | 1.35E+08 | 0.9995 | 1.15E-09 | 2.31E-09 |
| PHE | 6.22E+06 | 4.25E+08 | 0.9904 | 2.29E-09 | 4.59E-09 |
| Pro | 3.43E+06 | 7.34E+07 | 0.9982 | 5.34E-10 | 1.07E-09 |
| Ser | 1.22E+06 | 8.23E+07 | 0.9997 | 5.88E-10 | 1.18E-09 |
| Thr | 4.01E+06 | 2.77E+08 | 0.9989 | 7.31E-10 | 1.46E-09 |
| Try | 4.59E+05 | 3.12E+07 | 0.9996 | 4.47E-09 | 8.93E-09 |
| Tyr | 2.14E+06 | 1.47E+08 | 0.9993 | 2.43E-09 | 4.86E-09 |
| Val | 2.12E+07 | 9.58E+08 | 0.9714 | 1.04E-09 | 2.07E-09 |

**Table 3.**
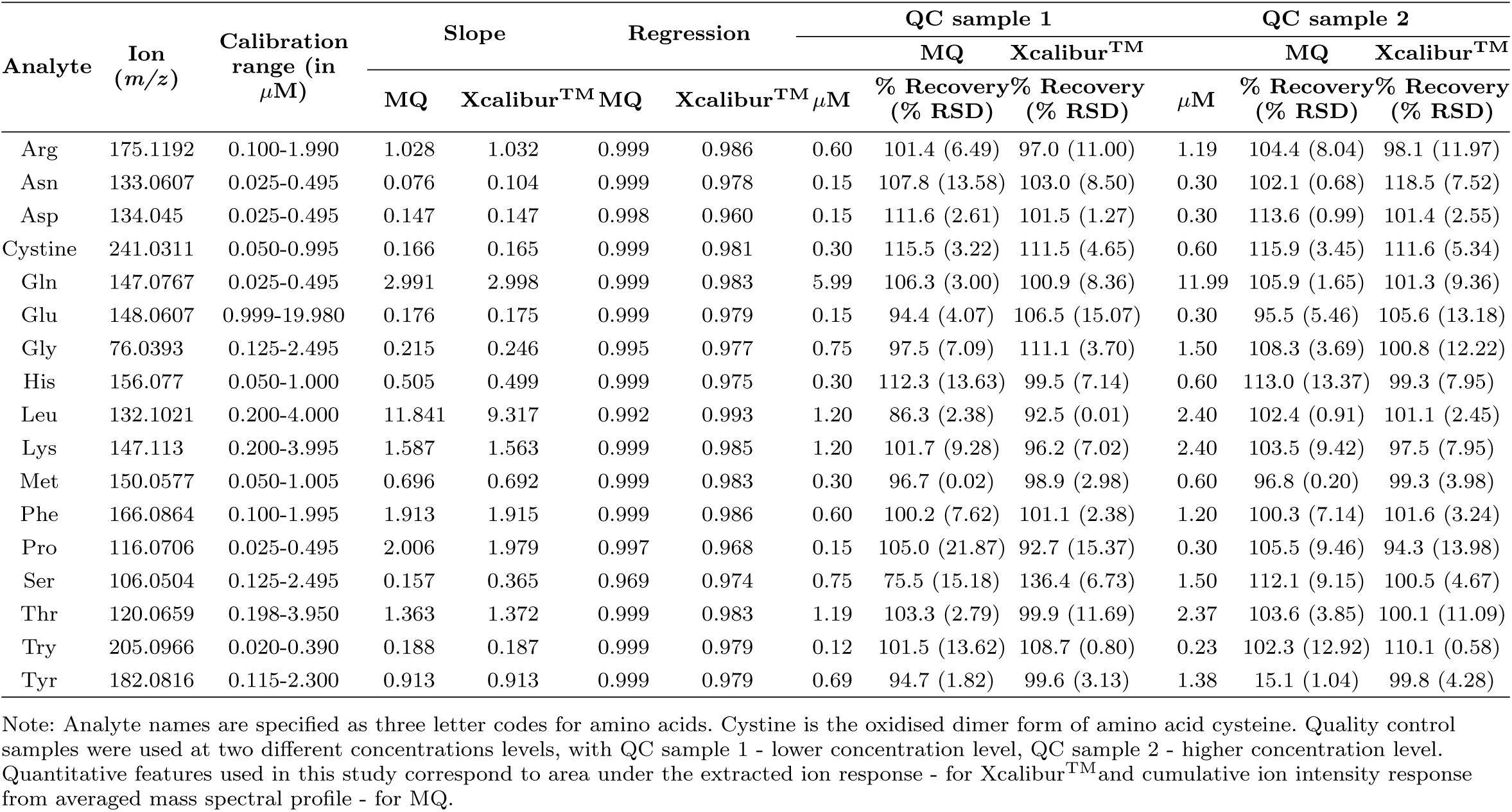
Quantitative information obtained subsequent to the LC-HRMS analysis of amino acids from a chemically defined mammalian cell culture media. Results obtained from MQ data processing were benchmarked using Xcalibur™(Thermo Fisher Scientific).

### 2.3 MALDI QTOF MS based quantitative analysis of biomarker metabolites

For generating MALDI Q-TOF MS data, a standard solution of SAM and SAH was prepared in methanol : water (1:1, v/v). For generating calibration curves, standard solutions were serially diluted to obtain the predetermined calibration levels and QCs as shown in Table 4. An internal standard (IS) solution consisting of 5.34 *µ*M melamine was uniformly spiked in all the samples before analysis to evaluate the performance and for data normalization. To carry out MALDI Q-TOF MS analysis, previously prepared standards were mixed in 1:1 ratio with 10 mg/ml of 2,5-DHB matrix solution, which was separately prepared in acetonitrile : 0.1% TFA in water (1:1, v/v). All samples were spotted on the MALDI target plate by dispensing 1 *µ*L of matrix-analyte mixture in 4 replicates for each of the calibrants. The data was acquired on Waters Synapt HDMS™MALDI Q-TOF instrument in positive ion mode.

**Table 4.** Quantitative information obtained from MQ data processing subsequent to MALDI-HRMS analysis of various analytes

| Analyte | Ion ($m/z$) | Calibration<br>range<br>(in $\mu\text{M}$) | Slope | $r^2$ | QC sample 1 | | QC sample 2 | |
| --- | --- | --- | --- | --- | --- | --- | --- | --- |
|  |  |  |  |  | % |  | % |  |
| | | | | | $\mu\text{M}$ | Recovery<br>(% RSD) | $\mu\text{M}$ | Recovery<br>(% RSD) |
| SAM<sup>a</sup> | 399.1395 | 0.75-10 | 0.054 | 0.97 | 2.5 | 78 (22.5) | 8 | 110 (10.4) |
| SAH<sup>a</sup> | 385.1286 | 0.75-10 | 0.237 | 0.93 | 2.5 | 73 (10.9) | 8 | 116 (12.2) |
| Atorvastatin<sup>b</sup> | 559.2603 | 1-10 | 0.089 | 0.96 | 3.5 | 100 (12.62) | 7.5 | 102 (5.3) |

Note: <sup>a</sup>Biomarker metabolites analysed using Waters Synapt HDMS<sup>TM</sup> MALDI Q-TOF.
<sup>b</sup>Pharmaceutical drug analysed using Q-Exactive (Thermo Fisher Scientific) equipped with AP-MALDI from Mass Tech Inc. USA.

### 2.4 Quantitative analysis of atorvastatin using AP-MALDI HRMS

Quantitative analysis of atorvastatin was performed on Q-Exactive (Thermo Scientific) mass spectrometer equipped with an atmospheric pressure matrix-assisted laser desorption ionization (AP-MALDI) source from Mass Tech Inc. USA. The instrument was operated in positive ion mode and the data was acquired within a mass range of 200-600 *m/z* at 70,000 FWHM resolution. 100 *µ*M stock solutions of atorvastatin were prepared in methanol : water (1:1, v/v). The stock solution was serially diluted to prepare predetermined calibration levels and QCs. 2, 5-DHB (10 mg ml^-1^) was prepared in acetonitrile : 0.1% TFA (1:1, v/v) and used as the MALDI matrix. 1.5 *µ*L of matrix was spotted on MALDI target plate. Samples premixed with 0.5 *µ*M verapamil, used as an internal standard, were subsequently spotted on dried matrix layer. Following the acquisition, the raw data was analyzed using MQ.

## 3 Overview of MQ

MQ features key aspects of data analysis as depicted in Figure 1 and includes a graphical user interface (GUI). Several modules within MQ have been designed to enable peak qualification, feature extraction, relative as well as absolute quantification, and untargeted analysis. Modules namely ‘Spectrum viewer’, ‘Isotopic confirmation’, ‘Quan calibration’, ‘Quan prediction’, ‘Relative quantitation’, ‘Database query’ and ‘Multivariate analysis - PCA’ are bundled into a single platform, as shown in Figure 2, to aid seamless user experience. An overview of HRMS data analysis workflow using MQ is illustrated in Figure 3. Various aspects of data processing using MQ are described below.

**Fig. 2.**
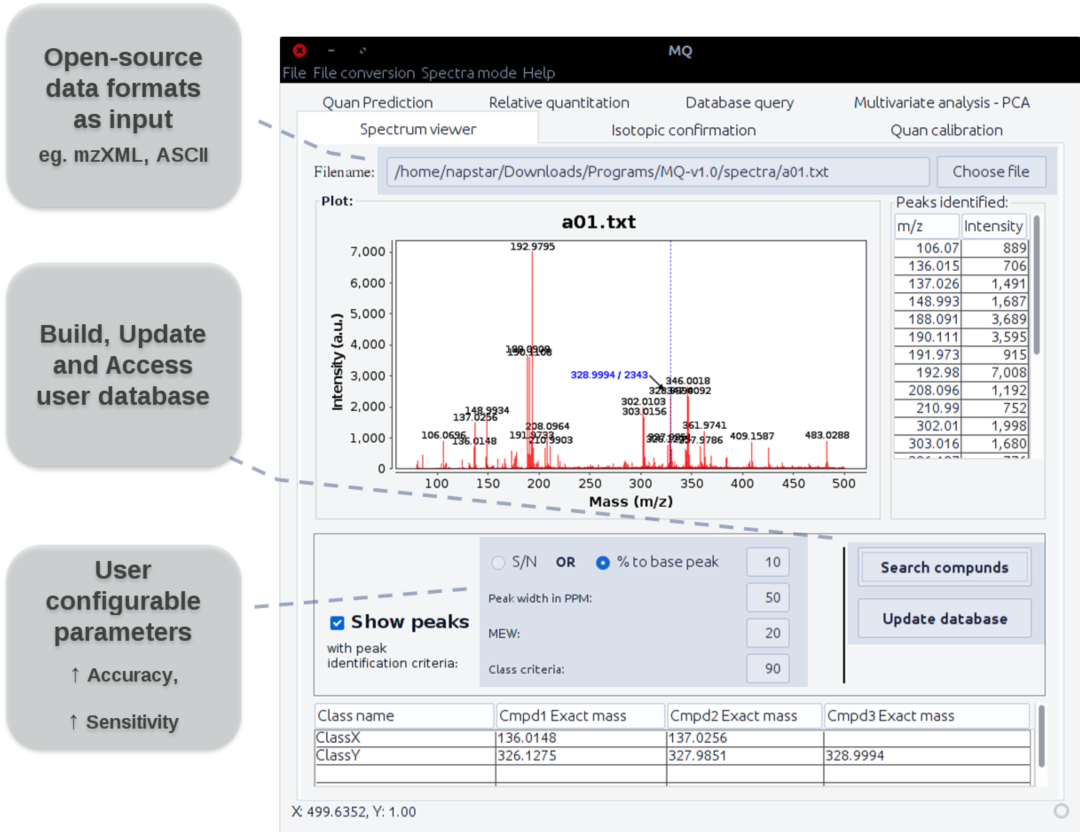
Screenshot for MQ user interface with available data analysis modules.

**Fig. 3.**
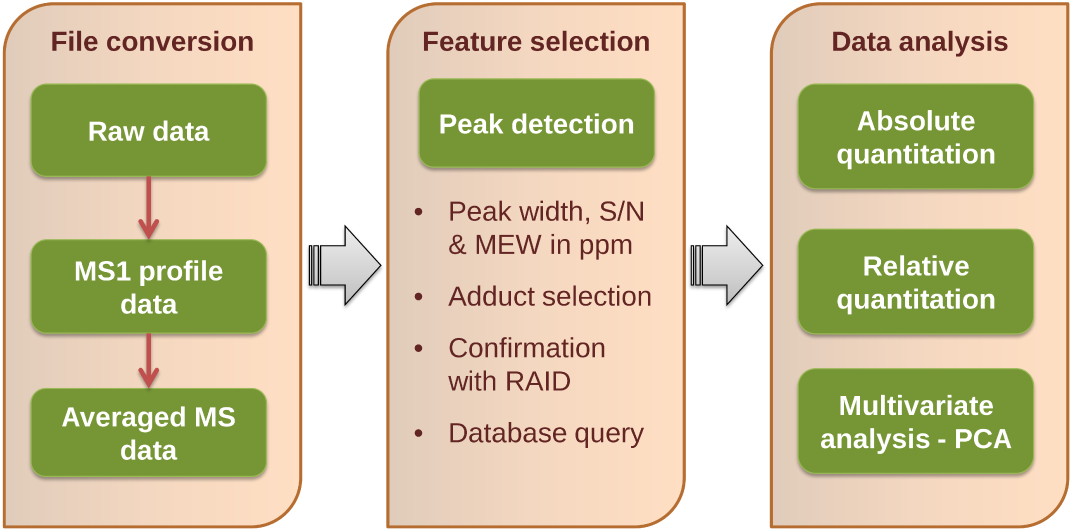
Schematic of HRMS data processing steps using MQ.

### 3.1 Data preprocessing

For data analysis with MQ, time averaged HRMS data is used as input data source in generic ASCII format (spectral *m/z*, intensity list) or mzXML format (common open source format developed by Seattle Proteome Center). Data analysis using time averaged MS spectrum serves as common platform for chromatography-based as well as direct MS-based approaches. LC-HRMS data can be considered as a stack of mass spectra acquired over a period of chromatographic run time. It contains details of ions detected during the process, generating significantly large datasets of information. In a typical LC-HRMS based quantitative analysis peak area, estimated as an extracted ion chromatogram (XIC), is used as a quantitative parameter. This is usually a response of ion current observed from an elution profile specific to an analyte of interest. For quantitative estimations using MQ, this multidimensional data (ion current over the *m/z* range as a function of LC runtime) is transformed into an averaged two dimensional mass spectral profile over a specific LC runtime. An averaged HRMS data profile was found to preserve the quantitative features specific to analyte peaks from sample. This approach was benchmarked against Xcalibur™(Thermo Scientific), which follows XIC based quantitative workflow.

Supporting module for generating average MS profile spectra from encrypted manufacturer specific file format is provided as an accompanying file conversion tool in MQ. File conversion using this tool is a two-step process. First the instrument specific file formats are converted to their respective MS1 (MS level 1) profiles using ‘MSConvert’ module from ProteoWizard package [7]. In the second stage, the MS1 profile data is averaged using in-house built tool ‘MSAvg’, written in Perl scripting language and GNU Octave environment, to generate two dimensional mass spectral profiles. Owing to the non-homogeneity in distribution of *m/z* positions across each scan, MSAvg identifies a list of unique *m/z* data entries from all mass spectral scans acquired in a chromatographic run. While averaging each scan, an interpolated spectral profile is generated using these unique *m/z* lists as input key for linear interpolation. For such interpolation, the spectral profile needs to be acquired in profile mode, which also imposes as a limitation for its inflexibility towards centroid data. Additionally, presence of isomeric analytes might also interfere with quantitative characteristics for respective analyte peak from averaged mass spectral profile as against chromatographic data. The final averaged mass spectral profile is used for all qualitative and quantitative analysis under different modules of MQ.

### 3.2 Spectrum viewer

Direct visual analysis of MS spectrum is often the quickest way for evaluating various qualitative checks such as presence/absence of a peak, mass accuracy (measured in ppm), signal intensity in ion counts, and the peak width. Spectrum viewer module of MQ offers direct visual analysis with additional options of optimizing peak finding criteria and subsequent database search. The spectrum viewer incorporates many user friendly and handy features such as, zoom-in/out, *m/z* value and intensity annotation. Additional options for spectra exporting (plot graphics to clipboard), saving spectra in image (png) format, and ‘properties’ option for improving visual attributes can also be availed. Users can also generate a list of peaks from the spectrum through optimization of ‘Peak finding criteria’ filters. Peaks can be qualified based upon signal-to-noise ratio (SNR), relative percentage intensity with respect to the highest or base peak, mass extraction window (MEW) within a set ppm and peak width. A convenient database search option is also available in the same window where annotated metabolite list or user created databases can be quickly referred. Additionally, in order to perform a high-throughput database query, a separate database search module is also available featuring similar ‘Peak finding criteria’ filters. Facility to add list of peaks generated from the spectrum viewer module as a user entry into database is possible.

### 3.3 Peak detection

Various methods for peak detection from MS data have been reported in last few decades. A recent review [9] provides an account on available feature extraction method along with their limitations. Few popular software tools such as, Mzmine [10] and XCMS [17] follows fitting of a template (Gaussian or Exponentially Modified Gaussian) function with given mass resolution setting for peak qualification. Various mass analyzers show varying response for ion distribution (resolution) at different *m/z* region [11]. This leads to difficulty in following template function based approach befitting for data from diverse list of MS instruments. MQ uses two step method for feature extraction. In the initial step, first derivative downward zero-crossing over method as peak picking algorithm with constraints over the slope threshold value is used [22]. First derivative spectra are subjected to moving average window (with width equivalent to half of MEW specified in ppm) smoothing. This helps in removing minor kinks that are a result of noise from the data, and increases computational efficiency in peak searching in such areas. Post smoothing first derivative spectra are subjected to peak finding for downward zero-crossing points, which essentially represents the highest point in a peak. Detected peaks are qualified based upon amplitude threshold and slope threshold, which keeps peak kurtosis in check. Slope threshold is defined based upon user specified peak width value as 0.5*(peak width points)^-2^, whereas amplitude threshold is based upon user defined filters such as S/N or relative percentage intensity to the base peak. Subsequently, in the second step a Gaussian function is fitted to the ion distribution observed in the proximity of peak. For this, a second order polynomial function is fit to log transformed ion count response for a set of points within a user defined MEW (in ppm) of peak that helps to capture Gaussian behavior of mass spectral profile. Peak area and peak intensity are estimated from this Gaussian function as area under the curve or based upon peak amplitude value estimation for polynomial fit, respectively.

### 3.4 Qualitative confirmation of analytes using RAID

Natural abundance for heavier isotopes of various elements leads to characteristic relative abundance for isotope intensity distribution (RAID) of analyte peaks, which is specific to elemental compositions. Various reports in past decade highlighted significance of RAID over and above mass accuracy offered by ultra-high resolution MS in characterization of analytes assertively [8, 14]. In case of complex and large molecules, prediction of RAID becomes more complicated and computationally intensive [1, 12]. In a recent publication, a web based tool for Molecular Isotopic Distribution Analysis (MIDAs) was developed with two improved algorithms for RAID calculation based on polynomial and Fourier-transform methods, having better performance in comparison to published tools for RAID estimation [1]. In MQ, an implementation of polynomial based algorithm from MIDAs web tool for fine grained RAID (used with high-resolution MS) estimation has been incorporated. Provision for adjustable mass accuracy and customizable isotopic abundances are salient features making it amenable to adapt for different experimental designs, such as isotopic labeling.

In brief, RAID estimation following polynomial based method involves multiplication of polynomial expressions for each element. These polynomial expressions are constituted from observed natural isotopic abundance values for each element. For fine grained RAID estimation, polynomial expansion was achieved following multinomial theorem with constraints over allowable exponent for individual isotope abundances in each element. These constraints are function of natural abundance of isotopes and elemental composition of analyte. Further details of algorithm can be found in article by Alves G. *et al.* [1].

With the provision of aforementioned algorithm in MQ for RAID estimation, screening and qualification of analyte with user specified elemental composition can be achieved from given mass spectrum. Confidence measure for qualification is provided as estimates of mass accuracy and percentage error in peak intensity for calculated heavier isotopic peak against observed peak.

### 3.5 Quantitative analysis

Quantitative analysis in MQ can be carried out based upon peak area or peak intensity. A weighted or non-weighted quantitative model with a linear/quadratic regression model fit can be generated for response curves extracted from peak areas or intensity values. Additionally, support for regression model fit for log transformed data is also offered. This can be used for adjusting varying instrumental responses and for fitting regression models of analytes whose concentration ranges vary over several orders of magnitude that usually affect their linear responses. Calibration models for absolute quantitation can be generated under ‘Quan-calibration’ module. Here, users can create and utilize a data library consisting of analytes and corresponding monoisotopic masses. An array of parameters can be specified for feature extraction and regression fit for list of analytes. These parameters are, analyte adduct ion(s), MEW (in ppm), weighted/non-weighted calibration model, *m/z* of internal standard used (optional), and spectral file names for the calibrants along with replicates. These parameters for a specific analysis project can be saved in a data library file. A least-square regression fit for all the analytes of interest is then generated that allows high-throughput simultaneous quantification in a single batch processing step. Adduct specific regression data is provided as an output. The best fitting adducts and models can then be selected based on regression statistics such as, slope, % RSD of technical replicates and intercept. Calibration models generated through ‘Quan-calibration’ module should be loaded in ‘Quan-prediction’ module to process the samples.

Relative quantification module allows direct comparison of internal standard normalized analyte responses across samples. Input parameters such as, *m/z* list for analyte ions, an internal standard adduct ion *m/z*, and MEW are required for processing. The output of relative quantitation provides the absolute and internal standard normalized peak area or intensity of analytes along with mass accuracy in ppm.

### 3.6 Untargeted profiling and multivariate analysis

MQ supports untargeted profiling and feature extraction using multivariate analysis in a metabolomics study. Data generated through full-scan HRMS has a multidimensional profile that poses a challenge for untargeted analysis. Unsupervised multivariate analysis is an unbiased means to identify a signature set of metabolites, which can be accounted as discriminative features. We have incorporated principal component analysis (PCA) as linear and unsupervised method for multivariate analysis of metabolomics data. PCA orthogonally transforms input spectral data by rotating the variables in coordinate space such that newly formed variables (Principle component factors-PC) should have maximum relevance with the variance within data. These transformations are a result of projecting original data points onto PC space identified by linear combination of the original variables, and thus it does not lead to loss of information. Additionally, a detailed analysis of these PC would help in identifying relevant list of analyte peaks, which holds higher coefficient values for linear combinations. These set of peaks are responsible for features represented by respective PC. For this multivariate analysis input variable data is provided either in terms of identified peak list with the use of ‘Peak finding criteria’ as mentioned in ‘Spectrum viewer’ module or intensity response for list of metabolites obtained by database query. Additional available set of parameters are: MEW for *m/z* grouping and normalization of peak intensity response using internal standard.

## 4 Results and Discussion

The performance of MQ was evaluated by simultaneous quantification of multiclass analytes that includes amino acids, metabolite biomarkers and pharmaceutical drug. The set of these three case studies was chosen to establish performance scaling for handling and analysis of data generated from varied analytical complexity, in addition to benchmarking using proprietary data analysis software.

A conventional chromatographic quantitative workflow for LC-HRMS analysis of a set of amino acids from biological matrix was used for comparison of data analysis using MQ and Xcalibur™(Thermo Scientific) software. Acquired data in native format with LC profiles was used as input for Xcalibur™based quantitative analysis, where peak area represented an area under curve of ion response observed from a chromatographic elution profile specific to an analyte of interest. In case of MQ, as discussed before, average mass spectral profile was used as input dataset for analysis. The accuracy and efficiency of quantitative workflows in MQ was evaluated at different levels of data processing such as, data conversion, identification of peak, peak integration, calibration curve fitting and unknown prediction. The calibration curves were examined in terms of %RSD of technical replicates, intercept, slope and regression coefficient (r^2^).

Evaluation of quantitative characteristic preserved by MSAvg, post data transformation of LC-HRMS data into two dimensional mass spectral profiles, is showcased in Table 2. A test sample representing a dilution level from calibration solutions of mammalian cell culture medium standard mixture was used for this comparison. Although the peak area estimated by Xcalibur™for LC-HRMS data was different in comparison to peak area, for respective list of analytes, estimated from averaged mass spectral profile using MQ, but a significant Pearsons correlation coefficient for a pairwise comparison was observed. This represents merits of preserved linear dependency of ion abundance for data transformed using MSAvg (Table 2). Further these peak areas were evaluated following sample test for equal variance (F-test). Significantly low estimated F-test statistic (in comparison to F-critical value of 0.0256) and p-value for rejection of null hypothesis, ratifies the unhindered quantitative nature of peak response following data transformation (Table 2).

Qualitative confirmation of all metabolites studied was evaluated using mass accuracy and natural abundance of heavier isotopes for each metabolite. Figure 4 shows screenshot for a sample spectra used for qualitative confirmation of glutamine using “Isotopic confirmation” module of MQ. Further, the output results of absolute quantitation of metabolites using MQ following LC-HRMS analysis is shown in Table 3. Results reported contain the comparison of slope and linear regression coefficients obtained for the calibrations processed using MQ and Xcalibur™. Regression coefficients of above 0.9 r^2^ indicate excellent linearity for the calibration curves within the used concentration range of 0.025 - 19.98 *µ*M, that are estimated in close proximity, by both MQ and Xcalibur™. The slopes for both the cases were also a close match largely and indicate similar responses and sensitivity for the methods. Two sets of QC samples to cover the broad calibration range were used to test the calibrations generated. The results obtained were reported as percentages relative to the expected recovery of 100%. The recoveries for all the amino acid QC samples for both the higher and lower concentration ranges were quite close to the expected recoveries and well within a generally acceptable precision of 15% relative standard deviation (%RSD). The data for proline shows greater deviation between MQ and Xcalibur™data. Most significantly, the recoveries obtained for the rest of the amino acid QC samples using MQ and Xcalibur™were strikingly similar. This is in spite of the fact that input for Xcalibur™was native raw data format, while for MQ it was transformed into averaged spectrum format. These results clearly establish that the spectral averaging preserves the quantitative characteristics of the data and benchmarks both qualitative and quantitative analysis using MQ.

**Fig. 4.**
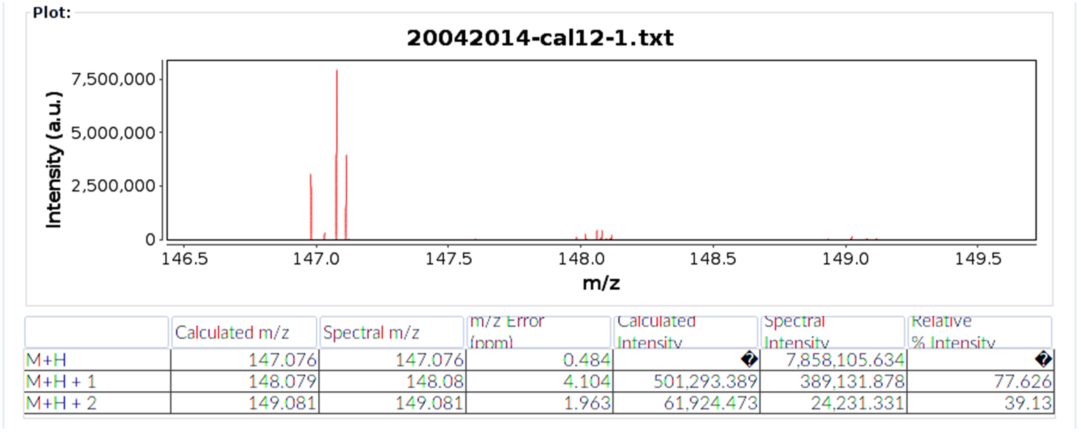
Screenshot for qualitative confirmation of glutamine using mass accuracy and RAID from a sample spectra

MQ was further tested successfully for the analysis of data acquired using chromatography free direct ionization source, MALDI coupled with TOF MS for metabolite disease biomarkers (S-adenosylmethionine: SAM, S-adenosylhomocysteine: SAH) and AP-MALDI coupled with Q-Exactive HRMS for pharmaceutical drug (Atorvastatin). SAM and SAH concentration levels are considered as a measure for cellular DNA methylation capacity and have been implicated in various pathological disorders [15, 5]. Whereas, atorvastatin is one of the most prescribed drugs belonging to the ‘statin’ class for treating high cholesterol levels. The calibration models were successfully generated and recoveries of QC samples were also estimated. Table 4 summarizes the quantitative information obtained from MQ data processing subsequent to data acquisition. The calibration ranges were within the concentration range of 0.5 to 10 *µ*M for SAM, SAH analysis and 1 to 10 *µ*M for atorvastatin. Calibration curves with excellent linearity were obtained with regression coefficient of above 0.9 r^2^ for all three analytes. Two QC samples, which cover higher and lower concentration range of calibration model, were used to estimate quantitative performance using measures of percentages recovery and percentage relative standard deviation for the estimations. For most of the QC samples % recovery obtained were closer to expected recovery concentration and with estimation precision below 15% in terms of %RSD. For SAM and SAH, % recovery estimated for lower concentration range QC was lower, especially with a higher deviation across replicates for SAM. This can be attributed to the spot-to-spot variations commonly observed in MALDI analysis, which can be improved using startegies like stable isotope labelling. Nonetheless, these results showcase the applicability of MQ for quantitative estimation of analytes using chromatography free direct ionization sources. Previously, MQ based quantitation of analytes from complex matrices such as plasma, urine and food matrix using MALDI based direct MS methods have also been reported.[2,16]

In addition, to showcase the specificity and sensitivity offered by MQ for feature selection and quantitative estimation of metabolites from complex biological matrix, extracellular milieu from a glioblastoma cancer cell line U87MG and its phenotypically different subpopulation of neurospheroidal (NSP) cell line was analyzed. LC-HRMS data of cell culture samples was used for comparative evaluation of relative quantitative estimations across the two cell lines (U87MG/NSP), for a set of amino acids, using relative quantitation module of MQ and ‘Quan browser’ module from proprietary software Xcalibur™. The results for this analysis are illustrated in Figure 5. As expected all the amino acids showed a comparable estimation of relative quantitative profiles, across MQ and Xcalibur™. These results further illustrates application of MQ for reliable quantitative analysis of metabolites from a complex biological matrix, with the help of time averaged mass spectral profile of LC-HRMS data.

**Fig. 5.**
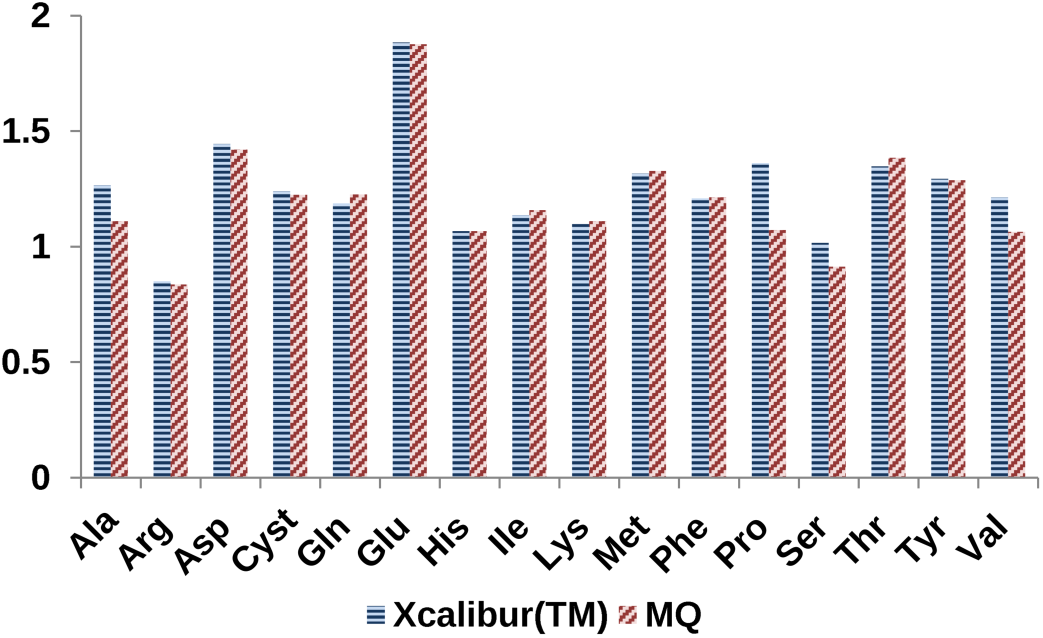
Quantitative performance of MQ in comparison to proprietary software Xcalibur™for a list of metabolites from cell culture samples. Analyte names are specified as three letter codes for amino acids. Cystine is the oxidised dimer form of amino acid cysteine.

## 5 Conclusion

Significant mass resolution offered by modern mass analyzers has encouraged the application of full scan mode MS analysis for reliable annotation of analytes along with qualitative and quantitative analysis. In order to accomplish this, a rigorous set of constraints that take into account high mass accuracies for peak qualification along with naturally present isotopic peak distributions are widely accepted criteria [8,14,11]. MQ incorporates the efficient MIDAs algorithm [1] for relative isotopic abundance confirmation towards this end. A high-throughput database query workflow and PCA based multivariate clustering analysis can further benefit qualitative elucidations. MQ offers flexibility with features such as, (a) availing mzXML and ASCII input data formats that are independent from proprietary raw data, (b) user configurable parameters for peak feature detection and (c) compatibility with both chromatography based and direct mass spectrometry methods. Seamless Qual-Quan integration is feasible using MQ through the benchmarked quantitative module that caters to both relative and absolute quantitation. Applications beyond the experimental scenarios showcased in this chapter are possible and include broad areas of food, pharmaceutical and clinical analysis.

## Acknowledgements

The authors thank Dr. Anu Raghunathan for providing mammalian cell culture medium solutions for LC-HRMS based quantitative analysis. The authors also thank Mylan Laboratories limited, Hyderabad (India) for providing atorvastatin standard sample. Dr. Ajeet Singh, Deepika Dhaware and Vishal Mahale for their valuable feedback in extensive evaluation of MQ.

## References

1. Alves, G., Ogurtsov, A.Y., Yu, Y.K.: Molecular Isotopic Distribution Analysis (MIDAs) with adjustable mass accuracy. J. Am. Soc. Mass Spectrom. 25(1), 57–70 (2014). DOI 10.1007/s13361-013-0733-7

2. Bhattacharya, N., Singh, A., Ghanate, A., Phadke, G., Parmar, D., Dhaware, D., Basak, T., Sengupta, S., Panchagnula, V.: Matrix-assisted laser desorption/ionization mass spectrometry analysis of dimethyl arginine isomers from urine. Anal. Methods 6(13), 4602–4609 (2014). DOI 10.1039/c4ay00309h

3. Hirosawa, M., Hoshida, M., Ishikawa, M., Toya, T.: MASCOT: multiple alignment system for protein sequences based on three-way dynamic programming. Comput. Appl. Biosci. 9(2), 161–167 (1993). DOI 10.1093/bioinformatics/9.2.161

4. Immanuel, S.R.C., Ghanate, A.D., Parmar, D.S., Marriage, F., Panchagnula, V., Day, P.J., Raghunathan, A.: Integrative analysis of rewired central metabolism in temozolomide resistant cells. Biochem. Biophys. Res. Commun. 495(2), 2010–2016 (2018). DOI 10.1016/j.bbrc.2017.12.073

5. James, S.J., Cutler, P., Melnyk, S., Jernigan, S., Janak, L., Gaylor, D.W., Neubrander, J.A.: Metabolic biomarkers of increased oxidative stress and impaired methylation capacity in children with autism. Am. J. Clin. Nutr. 80(6), 1611–1617 (2004). DOI 10.3945/ajcn.2008.26615.Am

6. Jellum, E., Kvittingen, E.A., Stokke, O.: Mass spectrometry in diagnosis of metabolic disorders. Biol. Mass Spectrom. 16(1-12), 57–62 (1988). DOI 10.1002/bms.1200160111

7. Kessner, D., Chambers, M., Burke, R., Agus, D., Mallick, P.: ProteoWizard: Open source software for rapid proteomics tools development. Bioinformatics 24(21), 2534–2536 (2008). DOI 10.1093/bioinformatics/btn323

8. Kind, T., Fiehn, O.: Metabolomic database annotations via query of elemental compositions: mass accuracy is insufficient even at less than 1 ppm. BMC Bioinformatics 7(1), 234 (2006). DOI 10.1186/1471-2105-7-234

9. Lytle, F.E., Julian, R.K.: Automatic Processing of Chromatograms in a High-Throughput Environment. Clin. Chem. 62(1), 144–53 (2016). DOI 10.1373/clinchem.2015.238816

10. Pluskal, T., Castillo, S., Villar-Briones, A., Oresic, M.: MZmine 2: modular framework for processing, visualizing, and analyzing mass spectrometry-based molecular profile data. BMC Bioinformatics 11(1), 395–405 (2010). DOI 10.1186/1471-2105-11-395

11. Rochat, B., Kottelat, E., McMullen, J.: The future key role of LChigh-resolution-MS analyses in clinical laboratories: a focus on quantification. Bioanalysis 4(24), 2939–2958 (2012). DOI 10.4155/bio.12.243

12. Rockwood, A.L., Haimi, P.: Efficient calculation of accurate masses of isotopic peaks. J. Am. Soc. Mass Spectrom. 17(3), 415–419 (2006). DOI 10.1016/j.jasms.2005.12.001

13. Römpp, A., Spengler, B.: Mass spectrometry imaging with high resolution in mass and space. Histochem. Cell Biol. 139(6), 759–783 (2013). DOI 10.1007/s00418-013-1097-6

14. Roussis, S.G., Proulx, R.: Reduction of chemical formulas from the isotopic peak distributions of high-resolution mass spectra. Anal. Chem. 75(6), 1470–1482 (2003). DOI 10.1021/ac020516w

15. Sibani, S., Melnyk, S., Pogribny, I.P., Wang, W., Hiou-Tim, F., Deng, L., Trasler, J., James, S.J., Rozen, R.: Studies of methionine cycle intermediates (SAM, SAH), DNA methylation and the impact of folate deficiency on tumor numbers in Min mice. Carcinogenesis 23(1), 61–65 (2002). DOI 10.1093/carcin/23.1.61

16. Singh, A., Panchagnula, V.: High throughput quantitative analysis of melamine and triazines by MALDI-TOF MS. Anal. Methods 3(10), 2360–2366 (2011). DOI 10.1039/c1ay05262d

17. Smith, C.A., Want, E.J., O’Maille, G., Abagyan, R., Siuzdak, G.: XCMS: processing mass spectrometry data for metabolite profiling using nonlinear peak alignment, matching, and identification. Anal. Chem. 78(3), 779–787 (2006). DOI 10.1021/ac051437y

18. Strohalm, M., Hassman, M., Košata, B., Kodíček, M.: mMass data miner: An open source alternative for mass spectrometric data analysis. Rapid Commun. Mass Spectrom. 22(6), 905–908 (2008). DOI 10.1002/rcm.3444

19. Strohalm, M., Kavan, D., Novák, P., Volný, M., Havlícek, V.: mMass 3: a cross-platform software environment for precise analysis of mass spectrometric data. Anal. Chem. 82(11), 4648–4651 (2010). DOI 10.1021/ac100818g

20. Sturm, M., Kohlbacher, O.: TOPPView: An open-source viewer for mass spectrometry data. J. Proteome Res. 8(7), 3760–3763 (2009). DOI 10.1021/pr900171m

21. Want, E.J., Masson, P., Michopoulos, F., Wilson, I.D., Theodoridis, G., Plumb, R.S., Shockcor, J., Loftus, N., Holmes, E., Nicholson, J.K.: Global metabolic profiling of animal and human tissues via UPLC-MS. Nat. Protoc. 8(1), 17–32 (2013). DOI 10.1038/nprot.2012.135

22. Yi, L., Dong, N., Yun, Y., Deng, B., Ren, D., Liu, S., Liang, Y.: Chemometric methods in data processing of mass spectrometry-based metabolomics: A review. Anal. Chim. Acta 914, 17–34 (2016). DOI 10.1016/j.aca.2016.02.001

23. Zhou, M., McDonald, J.F., Fernández, F.M.: Optimization of a direct analysis in real time/time-of-flight mass spectrometry method for rapid serum metabolomic fingerprinting. J. Am. Soc. Mass Spectrom. 21(1), 68–75 (2010). DOI 10.1016/j.jasms.2009.09.004

